# Towards a unifying framework for diversity and dissimilarity coefficients

**DOI:** 10.1101/2021.01.23.427893

**Authors:** Carlo Ricotta, László Szeidl, Sandrine Pavoine

## Abstract

Diversity and dissimilarity within and between species assemblages have now been studied for more than half a century by community ecologists in relation to their connections with ecosystem functioning. However, a generalized framework that puts diversity and dissimilarity coefficients under the same formal umbrella is still lacking. In this paper, we show that generalized means represent an effective tool to develop a unifying formulation for the construction of a large array of parametric diversity and dissimilarity measures. These measures include some of the classical diversity coefficients, such as the Shannon entropy, the Gini-Simpson index or the parametric diversity of Patil and Taillie, together with a large number of dissimilarity coefficients of the Bray-Curtis family and can be further extended to the measurement of functional and phylogenetic differences within and between plots.

## 1. Introduction

Diversity and dissimilarity measures have been typically used by community ecologists to explore the compositional heterogeneity within and between sample plots (or quadrats, sites, species assemblages, communities, etc.). For concave diversity measures, the usual way to link the within-plot heterogeneity (or *α*-diversity) of a pair of plots to their between-plot heterogeneity (*β*-diversity) is through Whittaker’s (1960) multiplicative decomposition. According to this model, *β*-diversity can be defined as the ratio between the pooled diversity of both plots (*γ*-diversity) and mean *α*-diversity in both plots such that 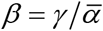.

Here, concavity means that the total diversity in the pooled pair of plots is not lower than the average diversity within each plot. From a mathematical viewpoint, let *U* and *V* be two plots where *x*_*Uj*_ and*x*_*Vj*_ are the (absolute) abundances of species *j* in plots *U* and 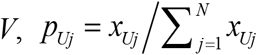 is the relative abundance of species *j* in plot *U* (with 0 ≤ *p*_*Uj*_ ≤ 1 and 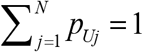), and *N* is the total number of species in both plots. Further, let *w*_*U*_ and *w*_*V*_ be two weights associated to plots *U* and *V* with 0 < *w*_*U*_ <1 and *w*_*V*_ = 1 − *w*_*U*_, and *γ* be the total diversity of the pooled pair of plots computed using the weighted mean of the species relative abundances in *U* and *V*: *p*_*γ j*_ = *w*_*U*_ *p*_*Uj*_ + *w*_*V*_ *p*_*Vj*_. For a concave diversity index, we have 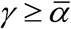 where 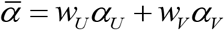 and *α*_*U*_ is the diversity of plot *U*. In its very essence, the requirement of concavity means that diversity increases by mixing.

According to this requirement, we thus have *β* ≥ 1, with *β* = 1 for two identical plots. Therefore, for concave diversity measures, the reciprocal of *β*-diversity 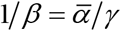 can be interpreted as an index of compositional similarity between *U* and *V* in the range [0,1]. Based on this approach and on its many successive improvements, a large number of diversity-related resemblance measures which depend on the excess of *γ*-diversity with respect to mean *α*-diversity have been defined (Rao 1982; Lande 1996; Jost 2007; Jost et al. 2010; Pavoine and Ricotta 2014; Chao et al. 2014, 2019).

Nonetheless, most of the ‘classical’ dissimilarity coefficients which have been used for decades to summarize compositional differences among plots, such as the Bray-Curtis dissimilarity, cannot be related to the difference of within-plot and between-plot diversity. Therefore, apart from Whittaker’s model, a generalized formulation that puts diversity and dissimilarity coefficients under the same umbrella is still lacking.

The aim of this paper is thus to propose a unifying framework for diversity and dissimilarity measures. This framework includes a large number of traditional dissimilarity coefficients of the ‘Bray-Curtis family’ and can be further extended to the measurement of functional and phylogenetic differences among plots. The paper is organized as follows: first, we provide an overview of a large class of diversity and dissimilarity measures, together with some of their properties. Next, we show that these measures can be traced back to a unified formulation which is based on the mathematical concept of generalized mean.

## 2. Diversity as the average rarity of a community

In their seminal paper, Patil and Taillie (1982) defined the diversity *D*_*U*_ of a given plot *U* as the average rarity of the species relative abundances *p*_*Uj*_:

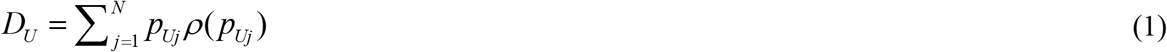

where the rarity *ρ* (*p*_*Uj*_) of species *j* is some decreasing function of its relative abundance *p*_*Uj*_. To avoid the additional complexity of species with zero abundances (particularly when dealing with effective numbers of species; see below), here and throughout the paper the summation is taken over all *N* species that are actually present in *U* with *p*_*Uj*_ > 0.

Diversity measures that fall within the definition of Patil and Taillie (1982) include two classical indices, such as the Shannon entropy (Shannon 1948) and the Simpson diversity (Simpson 1949). According to Shannon, the amount of information associated to a given plot *U* with species relative abundances *p*_*Uj*_ (*j* = 1, 2,…, *N*) can be summarized as:

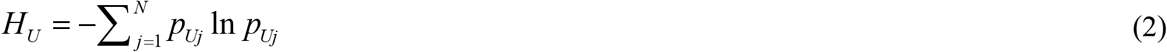

Where *ρ*(*p*_*Uj*_) = −ln(*p*_*Uj*_).

The Shannon entropy *H* is basically a measure of uncertainty in predicting the relative abundance of species in *U*. High entropy thus implies high unpredictability. Shannon’s entropy attains its maximum value *H*_*U*_ = ln *N* for a completely even community where all species occur in equal abundance (*p*_*Uj*_ = 1 *N*), whereas minimum entropy *H*_*U*_ = 0 is obtained if the community contains one dominant species whose relative abundance approaches unity and *N* − 1 species with vanishingly small abundances.

The amount of information obtained from observing the result of an experiment depending on chance can be taken to be numerically equal to the amount of uncertainty in the outcome of the experiment before carrying it out (Aczél and Daróczy 1975). Therefore, the Shannon entropy is usually interpreted as a measure of statistical information. Due to its logarithmic nature, a highly valued property of the Shannon entropy is additivity. That is, for two independent probability distributions *p* = (*p*_1_, *p*_2_, …, *p*_*N*_) and *q* = (*q*_1_, *q*_2_, …, *q*_*K*_) we have *H* (*pq*) = *H* (*p*) + *H* (*q*) where *pq* is the joint distribution of *p* and *q* (see e.g. Klir and Wierman 1999).

Since entropy reaches its maximum value when uncertainty is highest, information-theoretical measures appeared in early work on community structure. McArthur (1955) used the Shannon entropy to measure community stability, while Margalef (1958) first proposed to use entropy to summarize biological diversity. Since then, information-theoretical measures have become a fundamental tool for diversity analysis (Pielou 1966a, 1966b; He and Orlóci 1993).

The second pillar of classical diversity theory is the Simpson (or Gini-Simpson) index of diversity, which is defined as the probability that two individuals selected at random with replacement from a given community do not belong to the same species: 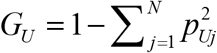

Apart from its probabilistic meaning, the Simpson index may be also formulated as an average community rarity:

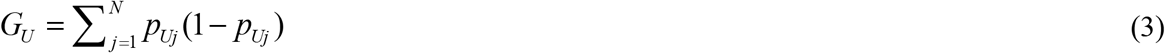

where the rarity function. *ρ*(*p*_*Uj*_) = (1 − *p*_*Uj*_) is linearly decreasing with the species relative abundances *p*_*Uj*_

## 3. Parametric Diversity

The Shannon and the Simpson indices are point descriptors of diversity. As such, they show only one portion of the whole diversity spectrum. To provide a vector description of diversity, the use of parametric functions has been advocated (Hurlbert 1971; Tóthmérész 1995). Rényi (1961) first introduced a parametric information measure, known as Rényi’s entropy, which is obtained by substituting the linear average in the Shannon entropy with the generalized Kolmogorov-Nagumo averages and by imposing the additivity of the information measures:

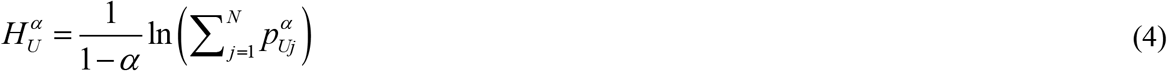

According to Eq. 4, there is a continuum of possible diversity measures that become increasingly dominated by the most abundant species for increasing values of the parameter α. The various diversity measures obtained by varying the parameter α are in fact different moments of the same basic function. The Rényi generalized entropy thus allows to represent community diversity by its diversity profile of 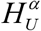 vs. α. As a result, rather than as a single-point summary statistic, parametric diversity can be seen as a scaling process that takes place in the topological data space (Podani 1992).

For 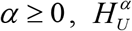 is concave. Therefore, the Rényi entropy is adequate to summarize community diversity in an ecologically meaningful manner. Due to its parametric nature,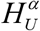 includes the three most important measures of species diversity: for 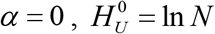 (i.e. a logarithmic transformation of species richness), for 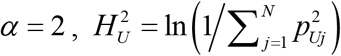, where 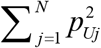 is the Simpson concentration or dominance (the opposite of diversity). For 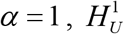 is undefined. However, its limit as α tends to unity gives the Shannon entropy: 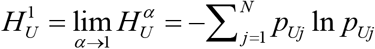.

Patil and Taillie (1982) further defined an additional parametric diversity index which has the form of an average community rarity as:

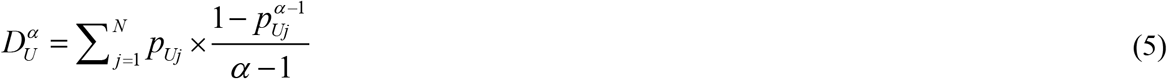

with 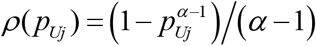.

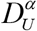 is a monotonic transformation of Rényi’s generalized entropy that has been used several times in many different fields (Havrdra and Charvát 1967; Daróczy 1970; Tsallis 1988; Keylock 2005). Like for Rényi’s entropy, 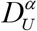 can generate each of the three classical diversity measures simply by varying the parameter α: for 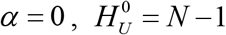 (i.e. a linear function of species richness that assigns zero diversity to single-species communities), for 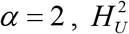 equals the Simpson index 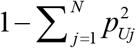, while for 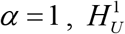 is defined in the limiting sense as the Shannon entropy 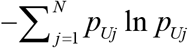. This led Lou Jost (2019) to say that parametric diversity functions were “a very interesting and important unification of what had once seemed like a smorgasbord of unrelated measures. It was the first sign that there might be a rich and deep «mathematics of diversity» that could bring order to the field”.

## 4. Diversity as effective numbers of species

McArthur (1965) and Whittaker (1972) noted that many diversity indices are nonlinear with respect to species addition. Therefore, even when all species are equally common, each added species leads to a smaller increment in overall diversity than the species added before it (see Jost et al. 2010). Hill (1973) suggested to solve this problem by converting traditional diversity measures to ‘effective numbers of species’. For a given diversity measure *D*_*U*_, the effective number of species or species equivalent *S*_*U*_, represents the number of equally abundant species (i.e. all with abundance *p*_*Uj*_ = 1 *S*_*U*_) that are needed in order that its diversity be *D*_*U*_.

Species equivalents are linear with respect to pooling, such that the species equivalent of a community of *N* totally dissimilar species in equal proportions is simply *N* (Leinster and Cobbold 2012). Jost (2006) further proved that the species equivalents of all measures of diversity that can be expressed as monotonic functions of 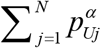, or limits of such functions as α approaches unity, are obtained as the exponential of the corresponding Rényi entropies of order α:

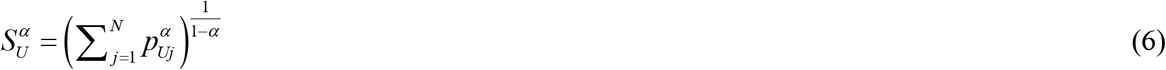

For details, see Jost (2006, Appendix 1). Hence, for 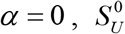 equals species richness *N*, for 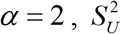 is the inverse of the Simpson concentration 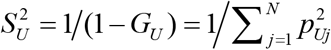 and for *α* = 1 the measure is defined in the limiting sense as the exponent of the Shannon entropy: 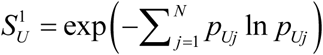. Effective numbers of species thus provide a general method for converting a wide variety of traditional diversity measures to the common currency of a species richness scale. For a throughout discussion on the properties of effective numbers of species, see Jost (2007) and Chao et al. (2014).

From Eq. 6 it follows that the effective numbers of species of all diversity functions that can be expressed as monotonic functions of 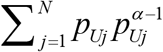, thus including all diversity functions of the form 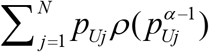, can be expressed as the reciprocal of a generalized mean of order *α* − 1 (Patil and Taillie 1982):

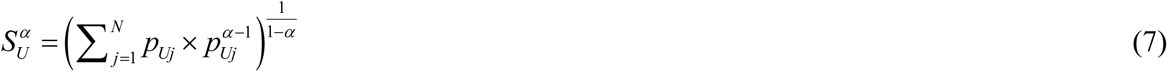

The generalized mean or power mean (Hardy et al. 1952) is a function that generalizes various notions of means, such as the arithmetic, or geometric mean into a single concept. Let *x*_1_, *x*_2_, …, *x*_*N*_ be a sequence of positive real values. For any weights *p*_*j*_ = (*p*_1_, *p*_2_, …, *p*_*N*_) adding up to 1, the classical arithmetic mean is 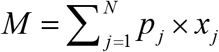. To generalize this notion, the power mean 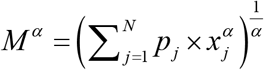 is obtained by transforming each value *x*_*j*_ into 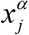, then by taking the weighted mean of the transformed values, and finally by applying the inverse transformation (Leinster and Cobbold 2012). According to this recipe, it is easily shown that Eq. 7 is the reciprocal of a generalized mean of order *α* − 1 of the species relative abundances *p*_*Uj*_ with weights *p*_*Uj*_.

Each generalized mean *M*^*α*^ always lies between the smallest and largest of the original values: min *x*_*j*_ ≤ *M*^*α*^ ≤ max *x*_*j*_. A few characteristic values of the parameter α recover more traditional notions of means. For example, for *α* = −∞, *M*^−∞^ = min *x*, for *α* = − 1, we obtain the harmonic mean 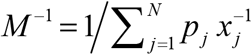, for *α* tending to 0 we obtain the geometric mean 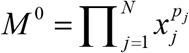, for *α* = 1 we obtain the classical arithmetic mean, while for *α* =∞, *M*^∞^ = max *x*_*j*_.

Consistently with our intuitive notion of diversity, for *α* ≥ 0, all moments of the parametric functions 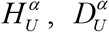 and 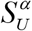 conform to Dalton’s (1920) principle of transfers, which states that diversity should increase if abundance is transferred from one species to another strictly less abundant species. This can be done either by transferring abundance to an already existing species, or by introducing a new species (i.e. by transferring abundance to a new species with zero abundance). Transferring abundance to an existing species increases community evenness, while introducing a new species increases richness (Patil and Taillie 1982).

## 5. From Simpson to Rao

All diversity measures discussed so far are based solely on the species relative abundances and cannot account for the ecological differences between species. Rao (1982) first introduced a diversity index, Quadratic diversity, that incorporates a measure of the pairwise differences between species. Quadratic diversity is defined as the expected dissimilarity between two individuals drawn randomly with replacement from the community:

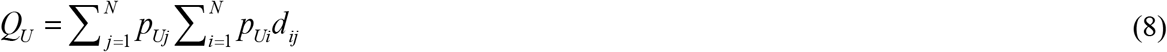

where *d*_*ij*_ is the dissimilarity among species *i* and *j* such that *d*_*ij*_ = *d*_*ji*_ and *d*_*jj*_ = 0.

These dissimilarities can be based either on functional or phylogenetic differences, as ecological differences between species are believed to be reflected in each of these. The mathematical properties of quadratic diversity have been extensively investigated by a number of authors (Shimatani 2001; Champely and Chessel 2002; Pavoine and Bonsall 2009; Rao 2010; Pavoine 2012) and the interested reader is addressed to these papers for details. Here, it is important to note that if the interspecies dissimilarities *d*_*ij*_ used to calculate the Rao index are squared Euclidean, *Q*_*U*_ is concave. A matrix of dissimilarities *d*_*ij*_ among *N* species is said to be squared Euclidean, if the *N* species can be embedded in a Euclidean space such that the Euclidean distance between species *i* and *j* is 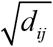 (Gower and Legendre 1986).

Assuming that biological differences between species are related to their dissimilarities, Leinster and Cobbold (2012) defined the ordinariness of species *j*, as the abundance of all species that are similar to *j* (including *j* itself):

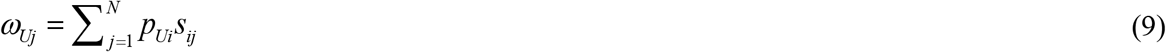

Where *s*_*ij*_ is the similarity between species *i* and *j* such that *s*_*ij*_ = 1 − *d*_*ij*_. Dealing with functional diversity, *ω*_*Uj*_ thus measures the commonness of all individuals in plot *U* that support the functions associated with species *j*. For 0 ≤ *d*_*ij*_ ≤ 1, *ω*_*Uj*_ ranges from *p*_*Uj*_ if all species *i* ≠ *j* are maximally dissimilar from *j*, to 1 if all species *i* ≠ *j* are functionally identical to *j*. Hence, *ω*_*Uj*_ ≥ *p*_*Uj*_.

Since 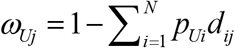, quadratic diversity can be thus expressed as the expected rarity of the species ordinariness *ω*_*Uj*_:

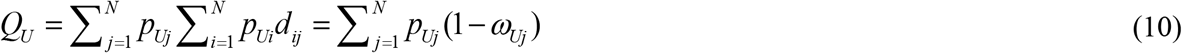

where the quantity 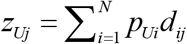 represents the abundance of all species in *U* that conflict with species *j* such that *z*_*Uj*_ + *ω*_*Uj*_ = 1.

According to Eq. 10, assuming that all species are maximally distinct from each other (i.e. if *d*_*ij*_ = 1 for all *i* ≠ *j*), *Q* reduces to the Simpson diversity 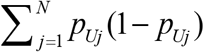. Rao’s *Q* can be thus interpreted as a measure of expected rarity among the species in a given assemblage if the species are not treated as maximally distinct from each other.

Expressing the generalized diversity of Patil and Taillie (1982) in terms of species ordinariness, Ricotta and Szeidl (2006) first obtained a parametric diversity function that incorporates a measure of pairwise differences between species:

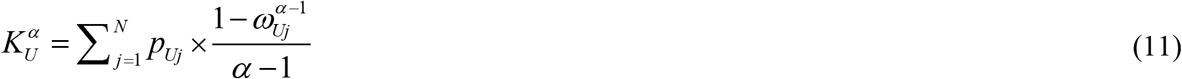

where, 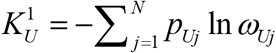 and 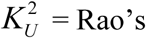 Rao’s quadratic diversity.

Note however that in the original paper of Ricotta and Szeidl (2006) there was an error in proving the concavity of 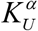 for *α* ≥ 2. In fact, using numerical simulations, it is possible to show that, apart for the special case *α* = 2, the parametric function 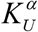 is generally not concave.

A few years later, Ricotta and Szeidl (2009) extended the concept of effective number of species to quadratic diversity. According to Ricotta and Szeidl (2009), the effective number of species of quadratic diversity is given by the inverse of Rao’s concentration 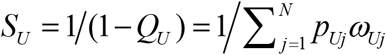. In this case, the effective number of species is defined as the number of equally abundant and *maximally dissimilar* species (i.e. with *d*_*ij*_ = 1 for all *i* ≠ *j*), needed to produce the given value of *Q*.

The effective number of species of the parametric diversity function of Ricotta and Szeidl 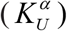 was then derived by Leinster and Cobbold (2012) as the reciprocal of a generalized mean of order *α* − 1 of the species ordinariness *ω*_*Uj*_ with weights *p*:

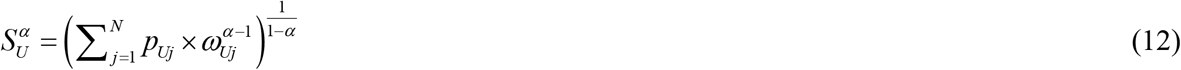

Like for all parametric diversity measures discussed so far, in Eq. 12, increasing the value of alpha increases the importance given to abundant or ordinary species compared to rare or distinct (unusual) species. Also, Eq. 11 and 12 satisfy an important condition discussed by Leinster and Cobbold (2012) and Botta-Dukát (2018) for functional and phylogenetic diversity measures which requires that diversity should not change if a given species *j* is replaced by two identical species with the same total abundance of *j*. For mathematical details, see Leinster and Cobbold (2012, Appendix A). A similar requirement was proposed by Weitzman (1992) and Solow and Polasky (1994) for the measurement of functional or phylogenetic richness, respectively, and by Pavoine and Ricotta (2019) for functional dissimilarity.

Diversity measures that conform to this branching requirement are generally not maximized if all species occur in equal abundance (*p*_*Uj*_ = 1 *N*). As a result, they do not conform to Dalton’s (1920) principle of transfers. Ricotta (2002) defined the indices that have their greatest value for non-completely even communities ‘weak diversity indices’ to differentiate them from ‘strong diversity indices’ such as the Shannon or the Simpson index, which show their maximum diversity for completely even communities. Although weak diversities are mathematically more difficult to handle, they are generally thought to describe ecological processes more convincingly that traditional diversities. While Poole (1974) defined classical diversity measures ‘answers to which questions have not yet been found’, there are now literally thousands of papers that relate functional or phylogenetic diversity to different aspects of community structure and species co-occurrence.

## 6. Anatomy of a dissimilarity index

Ecologists have developed a large number of measures to summarize the dissimilarity between a pair of plots (see e.g. Podani 2000; Legendre and Legendre 2012). Like for the Shannon or the Simpson diversity, most of these measures summarize community dissimilarity based either on species incidence or abundance data, thus assuming that all species are equally and maximally dissimilar from each other.

Irrespective of how dissimilarity is calculated, a desirable property for a dissimilarity coefficient is the so-called ‘sum property’. That is, its ability to be additively decomposed into species-level values. In this way, the practitioner can see which species contribute most to plot-to-plot dissimilarity (Ricotta and Podani 2017). For diversity indices, Patil and Taillie (1982) termed this property ‘dichotomy’ because the contribution of a given species to plot-to-plot diversity would be unchanged if the other species were grouped into a single complementary category.

Among the dissimilarity measures that conform to the sum property, the Bray and Curtis (1957) dissimilarity is obtained by standardizing the sum of species-wise differences by the total abundance of species in a pair of plots *U* and *V*:

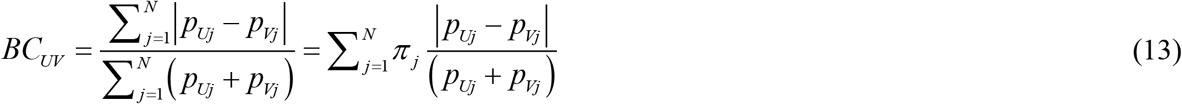

Where

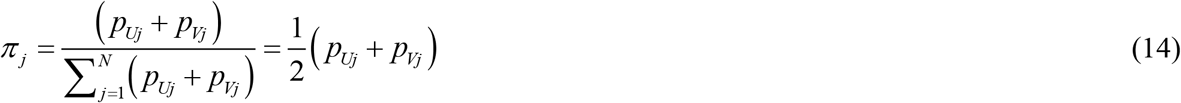

is the abundance of species *j* in *U* and *V* relative to the total species abundance in both plots. Note that, to keep a formal homogeneity with parametric diversity, in Eq. 13 the Bray-Curtis dissimilarity is expressed in terms of species relative abundances *p*_*Uj*_. Nonetheless, the same coefficient, together with all other dissimilarity coefficients used in this study, can be also expressed in terms of species absolute abundances *x*_*Uj*_ (see Ricotta and Pavoine 2015). Note also that, like for diversity measures, in Eq. 13 and throughout this paper, the summation is taken over all *N* species that are actually present in either *U* or *V* (i.e. *p*_*Uj*_ + *p*_*Vj*_ > 0).

Using the Bray-Curtis coefficient as a model, Pavoine and Ricotta (2019) proposed a general notation for the dissimilarity between two plots *U* and *V* that can be expressed as the mathematical expectation of the species-wise dissimilarity functions Δ(*p*_*Uj*_, *p*_*Vj*_) for all species in *U* and *V*.

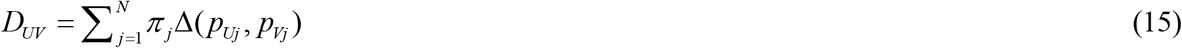

As shown in Eq. 13, the species-wise dissimilarity function of the Bray-Curtis coefficient can be formulated as Δ(*p*_*Uj*_, *p*_*Vj*_) = |*p*_*Uj*_ − *p*_*Vj*_| /(*p*_*Uj*_ + *p*_*Vj*_). However, other functions, such as the Marczewski-Steinhaus dissimilarity Δ(*p*_*Uj*_, *p*_*Vj*_) = |*p*_*Uj*_ − *p*_*Vj*_|/max{*p*_*Uj*_, *p*_*Vj*_} or any other dissimilarity function *sensu* Legendre (2014) with a numerator that summarizes the dissimilarity of species *j* between *U* and *V*, and a denominator that normalizes the index in the unit range may be equally used.

Another class of functions that conforms to Eq. 15 is composed of the many evenness-based dissimilarity coefficients proposed by Ricotta (2018) Δ(*p*_*Uj*_, *p*_*Vj*_) = 1 − *EVE*_*j*_, where *EVE*_*j*_ is a measure of the evenness of species *j* in plots *U* and *V*. For all these functions, the values of Δ(*p*_*Uj*_, *p*_*Vj*_) range from zero for two compositionally identical plots, to one for two maximally dissimilar plots. From what we learned in the previous paragraphs, it is now possible to generalize the dissimilarity function in Eq. 15 in two steps: first, by substituting the abundances *p*_*Uj*_ with the ordinariness *ω*_*Uj*_ in the species-wise dissimilarity function Δ (see Pavoine and Ricotta 2019), and next by substituting the arithmetic mean of Δ with the generalized mean:

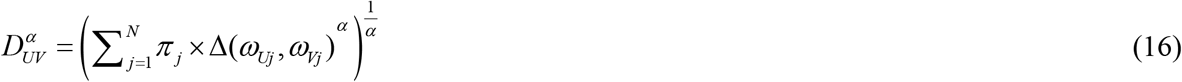

Eq. 16 represents a parametric formulation of plot-to-plot dissimilarity that includes pairwise differences between species. For *α* = 1, if all species in *U* and *V* are maximally dissimilar from each other, Eq. 16 reduces to Eq. 15, thus recovering all classical dissimilarity measures of the ‘Bray-Curtis family’. In addition, if all species are maximally dissimilar from each other and Δ(*p*_*Uj*_, *p*_*Vj*_) = *p*_*Uj*_ − *p*_*Vj*_ (*p*_*Uj*_ + *p*_*Vj*_), Eq. 16 can be expressed as a weighted version of the classical Minkowski distance 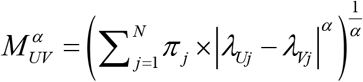 with *λ*_*Uj*_ = *p*_*Uj*_ (*p*_*Uj*_ + *p*_*Uj*_) and *λ*_*Vj*_ = *p*_*Vj*_ /(*p*_*Uj*_ + *p*_*Vj*_). Like for the Minkowski distance, in Eq. 16 the parameter alpha is related to the distinctness between plots, such that increasing the value of alpha increases the importance of large species-wise differences between plots compared to small differences.

It is easily shown that the generalized dissimilarity in Eq. 16 conforms to the branching requirement. According to this requirement, the functional dissimilarity between plots *U* and *V* does not change if one species in *U* or *V* is replaced by two identical species with the same total abundance. For details and proofs, see Pavoine and Ricotta (2019, Appendix 1).

## 7. Worked example

### 7.1 Methods

We used data on Alpine vegetation collected by Caccianiga et al. (2006). The data, which have been used in several papers for exploring community assembly rules along ecological gradients (e.g. Caccianiga et al. 2006; Ricotta et al. 2016; Ricotta et al. 2020) can be found in Ricotta et al. (2016, Appendix S2) and contain the abundances of 45 Alpine plant species collected in 59 plots of roughly 25 m^2^ in size along a primary succession on glacial deposits of the Rutor glacier (northern Italy). The species abundances in each plot were measured with a five-point ordinal scale transformed to ranks. The plots were then assigned to three successional stages based on the age of the glacial deposits: early successional stage (ESS; 17 plots), mid successional stage (MSS; 32 plots), and late successional stage (LSS; 10 plots). See Caccianiga et al. (2006) for additional details.

For each species, six quantitative traits were selected: canopy height (CH; mm), leaf dry matter content (LDMC; %), leaf dry weight (LDW; mg), specific leaf area (SLA; mm^2^ × mg^−1^), leaf nitrogen content (LNC; %), and leaf carbon content (LCC; %). Overall, these traits provide a good representation of the species global spectrum of form and function (Diaz et al. 2016). All data can be found in Caccianiga et al. (2006, pp. 16-17). All traits were standardized to zero mean and unit standard deviation. Based on the standardized trait values, we calculated a matrix of functional Euclidean distances *d*_*ij*_ between all 45 Alpine species. These distances were then rescaled in the range [0,1] by dividing each distance by the maximum value in the dataset.

First, we averaged the species abundances across the plots within each stage. Next, we calculated the species relative abundances within each stage. Finally, based on the species relative abundances, we calculated the parametric dissimilarity between all successional stages according to Eq. 16 with Δ(*ω*_*Uj*_, *ω*_*Vj*_) = |*ω*_*Uj*_ − *ω*_*Vj*_|**/** (*ω*_*Uj*_ + *ω*_*Vj*_). All calculations were done with a new R script available in Appendix 1. The reason for using the species relative abundances instead of absolute abundances for the calculation of functional dissimilarity is that we were interested in exploring how the ecological strategies of the vegetation change along the primary succession (i.e. how the functional characters are proportionally distributed among the species), irrespective of the species absolute abundances at each stage.

### 7.2 Results

According to previous work of Caccianiga et al. (2006) and Ricotta et al. (2020), the three successional stages are functionally well distinct. Along the primary succession, the vegetation is characterized by significant increase of leaf dry matter and leaf carbon content and a decrease of specific leaf area and leaf nitrogen content (see Ricotta et al. 2020). The functional dissimilarity profiles of 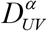 vs. α among the three successional stages of the Rutor glacier are shown in Figure 1. For all curves, functional dissimilarity tends to increase with increasing values of the parameter α according to a sigmoid pattern. For negative values of α, the curves show very low dissimilarity values. For values of α around zero, functional dissimilarity experiences a sharp growth. Finally, for increasingly positive values of α, functional dissimilarity increases asymptotically to its upper limit of max Δ(*ω*_*Uj*_, *ω*_*Vj*_).

**Figure 1.**
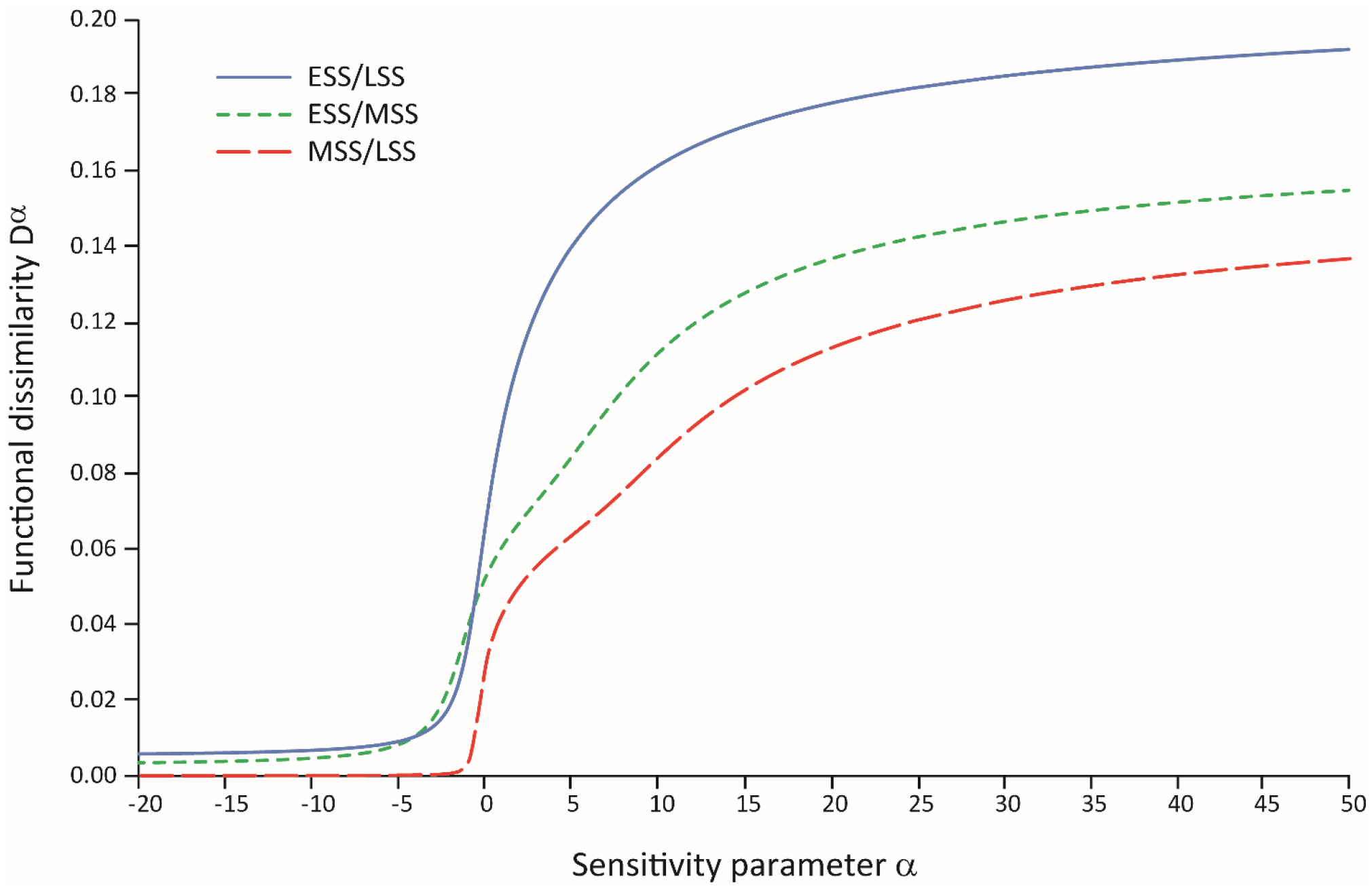
Dissimilarity profiles showing the functional differences among the three successional stages of the Alpine vegetation as a function of the sensitivity parameter α. ESS = early successional stage; MSS = mid successional stage; LSS = late successional stage.

Overall, the lowest functional differences are observed between the mid and the late successional stages, whereas, as expected, the highest differences are observed between the early and the late successional stages. These differences are particularly clear for positive values of α, thus emphasizing large species-wise differences between successional stages compared to small differences. In contrast, for negative values of α the curves of ESS vs. MSS and ESS vs. LSS cross, such that for *α* < 0 we cannot unambiguously say which of the two curves shows the largest functional differences.

## 8. Discussion

In this paper, we showed that generalized means represent an effective tool to develop a unifying notation for a large family of parametric diversity and dissimilarity functions. From a technical viewpoint, Chao et al. (2014) found that the diversity measures based on species ordinariness *ω*_*Uj*_ typically yield very low diversity values when the similarity matrix is computed from the species functional traits, and this causes problems for the interpretation. However, we suspect that this effect is due to the fact that in many cases the selected traits are too general and do not have any direct ecological association to the process of interest. Lavorel et al. (2008) emphasized that the traits that are actually relevant for ecosystem functioning depend case by case on the analyzed process. Therefore, a relevant aspect of functional analysis is the selection of an appropriate set of traits that optimize their association to the process of interest (Ricotta and Moretti 2010). We believe that advanced statistical methods, such as machine learning or artificial intelligence (Lucas et al. 2020) will greatly contribute to the construction of such ‘optimal’ functional spaces. In addition, like for many other summary statistics, for diversity and (dis)similarity indices, only ‘larger than’ comparisons may be valid. That is, the numerical values of those indices usually cannot be used as absolute indicators; they can only be used to rank the within- or between-plot heterogeneity in ways that are consistent with our ecological hypothesis (Kvålseth 2015).

From a more general perspective, in his comment on Patil and Taillie (1982), George Sugihara (1982) noted: “Truly ground-breaking contributions to the theory of species diversity are not likely to arise *in vitro* from a mathematical analysis of indices but will most probably depend on an interplay of analysis with real data”. A textbook example of this approach is the paper of Campbell Webb (2000) on the phylogenetic structure of ecological communities. Nonetheless, we think that addressing diversity theory in a more detached way may be equally beneficial, especially if our aim is to analyze the properties that diversity and dissimilarity measures should meet to summarize within- and between-plot heterogeneity in an ecologically appropriate manner.

Ecologists have developed a multitude of diversity and dissimilarity indices based on distinct goals and motivations. However, the choice of the most adequate index for solving a given ecological problem remains a complex question which does not have a clear and unequivocal answer (Ricotta and Podani 2017). Various properties have been advocated for diversity and dissimilarity indices; some of them are considered necessary by many authors, while others appear less relevant. However, given the multivariate nature of community data, the blanket is always too short and it is generally recognized that no single index can simultaneously possess even the most basic of these properties (Routledge 1983). For example, effective numbers of species usually do not possess the sum property. Therefore, in spite of their name, effective numbers of species cannot be decomposed into species-level patterns (Ricotta 2010).

Given this ‘uncertainty principle’, which is typical of most ecological indicators, we suggest to evaluate the index properties not (or not only) in abstract terms, but with reference to the needs of the specific questions at hand. For example, concavity which is generally considered an indispensable requirement for a diversity function, is really useful only if we want to calculate a measure of plot-to-plot community (dis)similarity that is based on the excess of *γ*-diversity with respect to mean *α*-diversity. Otherwise, if we focus solely on *α*-diversity, this requirement may no longer be needed. Note that, by relaxing the assumption that diversity needs to increase by mixing, we could also adopt a looser definition of concavity, for example by imposing that *γ* ≥ min{*α*_*U*_, *α*_*V*_} (see Avriel et al. 1988). Note also that the standard definition of concavity is based on an arithmetic mean, whereas some parametric diversity functions, such as all effective numbers of species in Eq. 7 and 12 are based on generalized means. Therefore, the standard definition of concavity does not necessarily fit these diversity functions perfectly (but see Routledge 1979; Chao et al. 2019).

A second important requisite for diversity and dissimilarity measures containing interspecies differences is that diversity/dissimilarity should not change if a given species *j* is replaced by two identical species with the same total abundance of *j* (Leinster and Cobbold 2012; Pavoine and Ricotta 2019). In functional terms, we can say that the measures that fulfill this branching requirement summarize the diversity/dissimilarity of ecosystem functions irrespective of the identity of the species that support them. In contrast, the measures that do not fulfill this requirement, such as Walker et al. (1999), Guiasu and Guiasu (2012), or Chiu and Chao (2014) to make only a few examples, basically summarize classical within- and between plot compositional heterogeneity calibrated by the functional or phylogenetic resemblance among species. Therefore, they are closer to classical measures of species diversity/dissimilarity than to their functional or phylogenetic analogues.

A final relevant requisite for diversity/dissimilarity measures is their ecological interpretability. According to Hurlbert (1971), the ecological meaning of the Shannon entropy and its parametric generalizations is at least dubious: “no one has yet specified exactly what significance the ‘number of bits per individual’ has to the individuals and populations in a community”. In addition, to the best of our knowledge, the foremost property of the Shannon entropy, additivity, has virtually never been used in modern community ecology and diversity theory. Therefore, the use of an overly complex logarithmic index without a clear ecological meaning is not necessarily the most appropriate choice for summarizing biological diversity.

On the other hand, the Simpson and the Rao diversity both have a clear and intuitive probabilistic meaning, the former in terms of interspecific encounters (see Hurlbert 1971; Patil and Taillie 1982), and the latter in terms of expected dissimilarity among species. Since probabilistic diversities are easier to interpret ecologically, a new class of parametric measures may be thus constructed by directly generalizing the Rao quadratic diversity. For example, Guiasu and Guiasu (2011) proposed to express the Rao quadratic diversity as a linear function of the joint distribution of the relative abundance of the species pairs. Let *π*_*ij*_ = *p*_*Ui*_ × *p*_*Uj*_ be the joint probability of the pair of species (*i, j*) in this order. Rao’s quadratic diversity can be thus rewritten as: 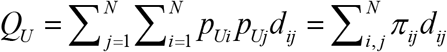 Accordingly, a natural way to generalize this function is by substituting the arithmetic mean with a power mean such that:

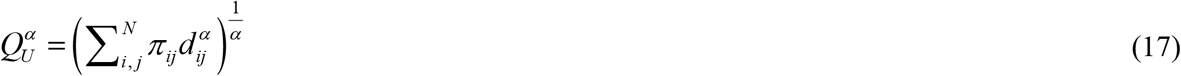

With *α* > 0. Eq. 17 thus represents the generalized mean of the pairwise interspecies distances *d*_*ij*_ weighted by the relative abundance of the corresponding species pairs. This index was first used by Rocchini et al. (2021) to summarize the diversity of remotely sensed primary productivity of the Earth’s terrestrial ecosystems. Although 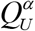 is generally not concave, except for the case *α* = 1, in Appendix 2 we will show that for 0 < *α* ≤ 1 it can be used to obtain a dissimilarity coefficient that is based on the excess of between-plot diversity with respect to mean within-plot diversity. Like for Eq. 16, increasing the value of the parameter alpha in Eq. 17 increases the importance of large differences between species, showing once again a direct relationship between parametric dissimilarity and diversity. In addition, like for all diversity measures discussed in this paper, for maximally dissimilar species, 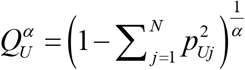 is maximized if all species occur in equal abundance.

This short example demonstrates that if we get out of the rather narrow cage of information-theoretical diversity and dissimilarity measures, there is virtually infinite space for new ‘targeted’ measures that are able to summarize within- and between-plot heterogeneity from many different viewpoints. In this paper, we focused mainly on generalized means, but this does not exhaust the range of available possibilities. Like any other ecological indicator, diversity and dissimilarity coefficients are part of a ‘complex, plural and dynamic approach to ecological studies’ (Juhász-Nagy 1984). Therefore, thinking about the existence of the ‘perfect index’ would simply be an illusion. Rather, a wide variety of imperfect measures are available, and their significance should be assessed based on their ability to address the specific ecological question under consideration. We hope, our present framework will contribute to transform this seemingly chaotic patchwork of unrelated diversity and dissimilarity measures into a coherent and structured system.

## Authors’ contributions

C.R. conceived the ideas and formulated the research problem. L.S. and S.P. provided important feedback which helped shape the research. C.R. and S.P. analyzed the data. C.R. took the lead in writing the main text of the manuscript, S.P. in writing the R script in Appendix 1, and L.S. in writing Appendix 2. All authors revised the manuscript critically and approved the final version.

## Supporting information

**Appendix 1**. R script for the calculation of parametric dissimilarity.

**Appendix 2**. Diversity partitioning of 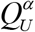.

## References

Aczél, J., Daróczy, Z. (1975) On measures of information and their characterizations. Academic Press, New York.

Avriel, M., Diewert, W.E., Schaible, S., Zang, I. (1988) Generalized Concavity. Plenum Press, New York.

Botta-Dukát, Z. (2018) The generalized replication principle and the partitioning of functional diversity into independent alpha and beta components. Ecography 41: 40–50.

Bray, J., Curtis, J. (1957) An ordination of the upland forest communities in southern Wisconsin. Ecological Monographs 27: 325–349.

Caccianiga, M., Luzzaro, A., Pierce, S., Ceriani, R.M., Cerabolini, B. (2006) The functional basis of a primary succession resolved by CSR classification. Oikos 112: 10–20.

Champely, S., Chessel, D. (2002) Measuring biological diversity using Euclidean metrics. Environmental and Ecological Statistics 9: 167–177.

Chao, A., Chiu, C.H., Jost, L. (2014) Unifying species diversity, phylogenetic diversity, functional diversity, and related similarity and differentiation measures through Hill numbers. Annual Review of Ecology, Evolution, and Systematics 45: 297–324.

Chao, A., Chiu, C.H., Wu, S.H., Wang, C.L., Lin, Y.C. (2019) Comparing two classes of alpha diversities and their corresponding beta and (dis)similarity measures, with an application to the Formosan sika deer Cervus nippon taiouanus reintroduction programme. Methods in Ecology and Evolution 10: 1286–1297.

Chiu, C.H., Chao, A. (2014) Distance-based functional diversity measures and their decomposition: a framework based on Hill numbers. PLOS ONE 9: e100014.

Dalton, H. (1920) Measurement of the inequality of incomes. Economic Journal 30: 348–361.

Daróczy, Z. (1970) Generalized information functions. Information and Control 16: 36–51.

Díaz, S., Kattge, J., Cornelissen, J.H.C., Wright, I.J., Lavorel, S. et al.. (2016) The global spectrum of plant form and function. Nature 529: 167–171.

Gower, J.C., Legendre, P. (1986) Metric and Euclidean properties of dissimilarity coefficients. Journal of Classification 3: 5–48.

Guiasu, R.C., Guiasu, S. (2011) The weighted quadratic index of biodiversity for pairs of species: a generalization of Rao’s index. Natural Science 3: 795–801.

Guiasu, R.C., Guiasu, S. (2012) The weighted Gini-Simpson index: revitalizing an old index of biodiversity. International Journal of Ecology 2012: 478728.

Hardy, G., Littlewood, J.E., Pólya, G. (1952) Inequalities. Cambridge University Press, Cambridge.

Havrda, M., Charvat, F. (1967) Quantification method of classification processes: concept of structural α-entropy. Kybernetica 3: 30–35.

He, X.S., Orlóci, L. (1993) Comparative diversity analysis of vegetation. Abstracta Botanica 17: 79–86.

Hill, M. (1973) Diversity and evenness: A unifying notation and its consequences. Ecology 54: 427–432.

Hill, M.O. (1973) Diversity and evenness: a unifying notation and its consequences. Ecology 54: 427–431.

Hurlbert, S.H. (1971) The nonconcept of species diversity: a critique and alternative parameters. Ecology 52: 577–586.

Jost, L. (2006) Entropy and diversity. Oikos 113: 363–375.

Jost, L. (2007) Partitioning diversity into independent alpha and beta components. Ecology 88: 2427–2439.

Jost, L. (2019) What do we mean by diversity? The path towards quantification. Mètode Science Studies Journal 9: 55–61.

Jost, L., DeVries, P.J., Walla, T., Greeney, H., Chao, A., Ricotta, C. (2010) Partitioning diversity for conservation analyses. Diversity and Distributions 16: 65–76.

Juhász-Nagy, P. (1984) Spatial dependence of plant populations. Part 2. A family of new models. Acta Botanica Hungarica 30: 363–402.

Keylock, C. (2005) Simpson diversity and the Shannon-Wiener index as special cases of a generalized entropy. Oikos 109: 203–207.

Klir, G.J., Wierman, M.J. (1999) Uncertainty-Based Information. Physica-Verlag, Heidelberg.

Kvålseth, T.O. (2015) Evenness indices once again: critical analysis of properties. SpringerPlus 4: 232.

Lande, R. (1996) Statistics and partitioning of species diversity, and similarity among multiple communities. Oikos 76: 5–13.

Lavorel, S., Grigulis, K., McIntyre, S., Williams, N.S.G., Garden, D. et al.. (2008) Assessing functional diversity in the field – methodology matters! Functional Ecology 22: 134–147.

Legendre, P., Legendre, L. (2012) Numerical Ecology. Elsevier, Amsterdam.

Leinster, T., Cobbold, C.A. (2012) Measuring diversity: the importance of species similarity. Ecology 93: 477–489.

Lucas, T.C.D. (2020) A translucent box: interpretable machine learning in ecology. Ecological Monographs 90: e01422.

MacArthur, R. (1965) Patterns of species diversity. Biological Reviews 40: 510–533.

MacArthur, R.H. (1955) Fluctuation of animal populations and a measure of community stability. Ecology 36: 533–536.

Margalef, D.R. (1958) Information theory in ecology. General Systems 3: 36–71.

Patil, G.P., Taillie, C. 1982. Diversity as a concept and its measurement. Journal of the American Statistical Association 77: 548–561.

Pavoine, S. (2012) Clarifying and developing analyses of biodiversity: towards a generalisation of current approaches. Methods in Ecology and Evolution 3: 509–518.

Pavoine, S., Bonsall, M.B. (2009) Biological diversity: distinct distributions can lead to the maximization of Rao’s quadratic entropy. Theoretical Population Biology 75: 153–163.

Pavoine, S., Ricotta, C. (2014) Functional and phylogenetic similarity among communities. Methods in Ecology and Evolution 5: 666–675.

Pavoine, S., Ricotta, C. (2019) Measuring functional dissimilarity among plots: Adapting old methods to new questions. Ecological Indicators 97: 67–72.

Pielou, E.C. (1966a) Shannon’s formula as a measure of species diversity: its use and misuse. American Naturalist 100: 463–465.

Pielou, E.C. (1966b) The measurement of diversity in different types of biological collections. Journal of Theoretical Biology 13: 131–144.

Podani, J. (1992) Space series analysis: processes reconsidered. Abstracta Botanica 16: 25–29.

Podani, J. (2000) Introduction to the Exploration of Multivariate Biological Data. Backhuys, Leiden, NL.

Poole, R.W. (1974) An Introduction to Quantitative Ecology. McGraw-Hill, New York.

Rao, C.R. (1982) Diversity and dissimilarity measurements: A unified approach. Theoretical Population Biology 21: 24–43.

Rao, C.R. (2010) Quadratic entropy and analysis of diversity. Sankhya 72: 70–80.

Rényi, A. (1961) On measures of information and entropy. Proceedings of the fourth Berkeley Symposium on Mathematics, Statistics and Probability 1960, Vol. 1. University of California Press, Berkeley, CA, pp. 547–561.

Ricotta, C. (2002) Bridging the gap between ecological diversity indices and measures of biodiversity with Shannon’s entropy: comment to Izsák and Papp. Ecological Modelling 152: 1–3.

Ricotta, C. (2010) On beta diversity decomposition: Trouble shared is not trouble halved. Ecology 91: 1981–1983.

Ricotta, C. (2018) A family of (dis)similarity measures based on evenness and its relationship with beta diversity. Ecological Complexity 34: 69–73.

Ricotta, C., Acosta, A.T.R., Caccianiga, M., Cerabolini, B.E.L., Godefroid, S., Carboni, M. (2020) From abundance-based to functional-based indicator species. Ecological Indicators 118: 106761.

Ricotta, C., de Bello, F., Moretti, M., Caccianiga, M., Cerabolini, B., Pavoine, S. (2016) Measuring the functional redundancy of biological communities: a quantitative guide, Methods in Ecology and Evolution 7: 1386–1395.

Ricotta, C., Moretti, M. (2010) Assessing the functional turnover of species assemblages with tailored dissimilarity matrices. Oikos 119: 1089–1098.

Ricotta, C., Pavoine, S. (2015) Measuring similarity among plots including similarity among species: an extension of traditional approaches. Journal of Vegetation Science 26: 1061–1067.

Ricotta, C., Podani, J. (2017) On some properties of the Bray-Curtis dissimilarity and their ecological meaning. Ecological Complexity 31: 201–205.

Ricotta, C., Szeidl, L. (2006) Towards a unifying approach to diversity measures: Bridging the gap between the Shannon entropy and Rao’s quadratic index. Theoretical Population Biology 70: 237–243.

Ricotta, C., Szeidl, L. (2009) Diversity partitioning of Rao’s quadratic entropy. Theoretical Population Biology 76: 299–302.

Rocchini, D., Marcantonio, M., Da Re, D., Bacaro, G., Feoli, E. et al.. (2021) From zero to infinity: minimum to maximum diversity of the planet by spatio-parametric Rao’s quadratic entropy. Global Ecology and Biogeography, doi: 10.1111/geb.13270.

Routledge, R. (1979) Diversity indices: Which ones are admissible? Journal of Theoretical Biology. 76: 503–515.

Routledge, R.D. (1983) Evenness indices: are any admissible? Oikos 40: 149–151.

Shannon, C. (1948) A mathematical theory of communication. Bell System Technical Journal 27: 379–423.

Shimatani, K. (2001) On the measurement of species diversity incorporating species differences. Oikos 93: 135–147.

Simpson, E.H. (1949) Measurement of diversity. Nature 163: 688.

Solow, A. R. & Polasky, S. 1994 Measuring biological diversity. Environmental and Ecological Statistics 1: 95–107.

Sugihara, G. (1982) Comment on Diversity as a concept and its measurement. Journal of the American Statistical Association 77: 564–565.

Tóthmérész, B. (1995) Comparison of different methods for diversity ordering. Journal of Vegetation Science 6, 283–290.

Tsallis, C. (1988). Possible generalization of Boltzmann-Gibbs statistics. Journal of Statistical Physics 52: 479–487.

Walker, B., Kinzig, A., Langridge, J. (1999) Plant attribute diversity, resilience, and ecosystem function: the nature and significance of dominant and minor species. Ecosystems 2: 95–113.

Webb, C.O. (2000) Exploring the phylogenetic structure of ecological communities: an example for rain forest trees. American Naturalist 156: 145–155.

Weitzman, M.L. (1992) On diversity. Quarterly Journal of Economics 107: 363–405.

Whittaker, R. (1972) Evolution and measurement of species diversity. Taxon 21: 213–251.

Whittaker, R.H. (1960) Vegetation of the Siskiyou Mountains, Oregon and California. Ecological Monographs 30: 279–338.

