## Appendix 1 for "Towards a unifying framework for diversity and dissimilarity coefficients"

**Appendix 1.** R script for the calculation of parametric dissimilarity.

To calculate parametric dissimilarity, we modified the function `generalized_Tradidiss` of package `adiv` (Pavoine 2020). For a full description of the function, see the manual of package `adiv`.

In this new version of the function, named `generalized_Tradidiss_alpha`, we added a new numeric parameter that corresponds to the parameter alpha of Equation 16. This updated version of the function will be included in `adiv` after the publication of the paper.

```
generalized_Tradidiss_alpha <-
function(comm, dis, method = c("GC", "MS", "PE"), abundance =
c("relative", "absolute", "none"), weights = c("uneven", "even"),
alpha = 1, tol = 1e-8)
{
    q <- alpha[1]
    if(!is.numeric(q)) stop("q must be a numeric")
    if(inherits(dis, "dist")) dis <- as.matrix(dis)
    if(any(dis>1)){
        warning("Phylogenetic dissimilarities are not in the range
0-1. They have been normalized by the maximum")
        dis <- dis/max(dis)
    }
    if(any(!colnames(comm) %in%rownames(dis))) stop("At least one
species in the matrix of abundances is missing in the matrix of
dissimilarities")
    if(any(!colnames(comm) %in%colnames(dis))) stop("At least one
species in the matrix of abundances is missing in the matrix of
dissimilarities")
    method <- method[1]
    if(!method%in%c("GC", "MS", "PE")) stop("Incorrect definition
for method")
    abundance <- abundance[1]
    if(!abundance%in%c("relative", "absolute", "none"))
stop("Incorrect definition for abundance")
    weights <- weights[1]
    if(!weights%in%c("uneven", "even")) stop("Incorrect definition
for weights")
dis <- dis[colnames(comm), colnames(comm)]
    dataset <- t(comm)
    similarities <- 1-as.matrix(dis)
total <- colSums(dataset)
    if(abundance == "relative")
        abu <- sweep(dataset, 2, total, "/")
    else if(abundance == "none"){
abu <- dataset
        abu[abu>tol] <- 1
        abu[abu<=tol] <- 0
}
    else abu <- dataset
    num.plot<-dim(dataset)[2]
    num.sp <- dim(dataset)[1]
    names<-list(colnames(dataset), colnames(dataset))
```

```
    dis.matrix<-matrix(0, nrow=num.plot, ncol=num.plot,
dimnames=names)
    for (i in 2:num.plot) {
    for (j in 1:(i-1)) {
        mat_folk <- similarities*abu[, j]
        mat_folk2 <- similarities*abu[, i]
        Zik <- colSums(mat_folk)
        Zih <- colSums(mat_folk2)
        tabZ <- rbind.data.frame(Zik, Zih)
        garde <- apply(tabZ, 2, sum)>tol & apply(abu[,c(i,j)], 1,
sum)>tol
Zik <- Zik[garde]
        Zih <- Zih[garde]
tabZ <- tabZ[, garde]
        if(weights == "even") wk <- rep(1/length(Zih), length(Zih))
        else {
        wk <- (abu[, j]+abu[, i])/sum(abu[, c(i,j)])
wk <- wk[garde]
        }
        if(method == "GC"){
            if(abs(q)<tol) index <- prod((abs(Zik-Zih)/apply(tabZ,
2, sum))^wk)
            else index <- (sum((wk*(abs(Zik-Zih)/apply(tabZ, 2,
sum))^q)))^(1/q)
        }
        else if(method == "MS"){
            if(abs(q)<tol) index <- prod((abs(Zik-Zih)/apply(tabZ,
2, max))^wk)
            else index <- (sum((wk*(abs(Zik-Zih)/apply(tabZ, 2,
max))^q)))^(1/q)
        }
        else {
            zik <- Zik / (Zik + Zih)
            zih <- Zih / (Zik + Zih)
            tabz <- rbind.data.frame(zik, zih)
            enlev <- as.vector(apply(tabz, 2,
                function(x) any(x<=tol)))
            zik <- zik[!enlev]
            zih <- zih[!enlev]
            wkN <- wk[!enlev]
            wkY <- wk[enlev]
            if(abs(q)<tol)
                index <- prod((1
                +(zik * log(zik, base = 2)
                + zih * log(zih, base = 2)))^wkN)
            else
                index <- (sum(wkY) + sum(wkN*(1
                +(zik * log(zik, base = 2)
                + zih * log(zih, base = 2)))^q))^(1/q)
        }
        dis.matrix[i, j] <- index
}
    }
    dis.matrix <- dis.matrix + t(dis.matrix)
    dis.matrix <- dis.matrix + diag(rep(0, num.plot))
return(as.dist(dis.matrix))
}
```

### Example of use

```
# The script below can be used to build the Figure 1 of the main text.

# First load the function generalized_Tradidiss_alpha in your R console

# Then use the following scripts

library(adiv) # this loads package adiv

data(RutorGlacier)

# this loads the dataset that is available in adiv


ab <- RutorGlacier$Abund  # Species abundances in plots

abmean <- apply(ab, 2, function(x) tapply(x, RutorGlacier$Fac,
mean))  # Average species abundances for each successional stage

traits <- RutorGlacier$Traits2[1:6] # Table of species traits

dis <- dist(scale(traits))  # Functional distances between species

dis<-dis/max(dis) # Functional distances normalized between 0 and 1

E <- seq(-20, 50, le=71) # values for parameter alpha

GCdis <- lapply(E, function(k) generalized_Tradidiss_alpha(abmean,
dis, method="GC", alpha=k)) # Calculation of the plot-to-plot
distances

# Note that the function may be time consuming with large datasets.

# Plot-to-plot distances are put below in a table

TGCdis <- cbind.data.frame(lapply(GCdis, as.vector))

colnames(TGCdis) <- paste("i", 1:71, sep="")

rownames(TGCdis) <- c("ESS/LSS", "ESS/MSS", "MSS/LSS")


# In table TGCdis, columns represent different values of alpha, ESS/MSS means comparison between early-successional stage and mid-successional stage, ESS/LSS means comparison between early-successional stage and late-successional stage, and MSS/LSS means comparison between mid-successional stage and late-successional stage


# The instructions below plot the results, which leads to Figure 1
of the main text:

plot(E, TGCdis[1, ], type="l", col="blue", xlab="Sensitivity
parameter alpha", ylab="Functional dissimilarity Dalpha")
```

```
points(E, TGCdis[2, ], type="l", col="green", lty=2)

points(E, TGCdis[3, ], type="l", col="red", lty=3)

legend("topleft", legend = rownames(TGCdis), lty=1:3,
col=c("blue","green","red"))
```

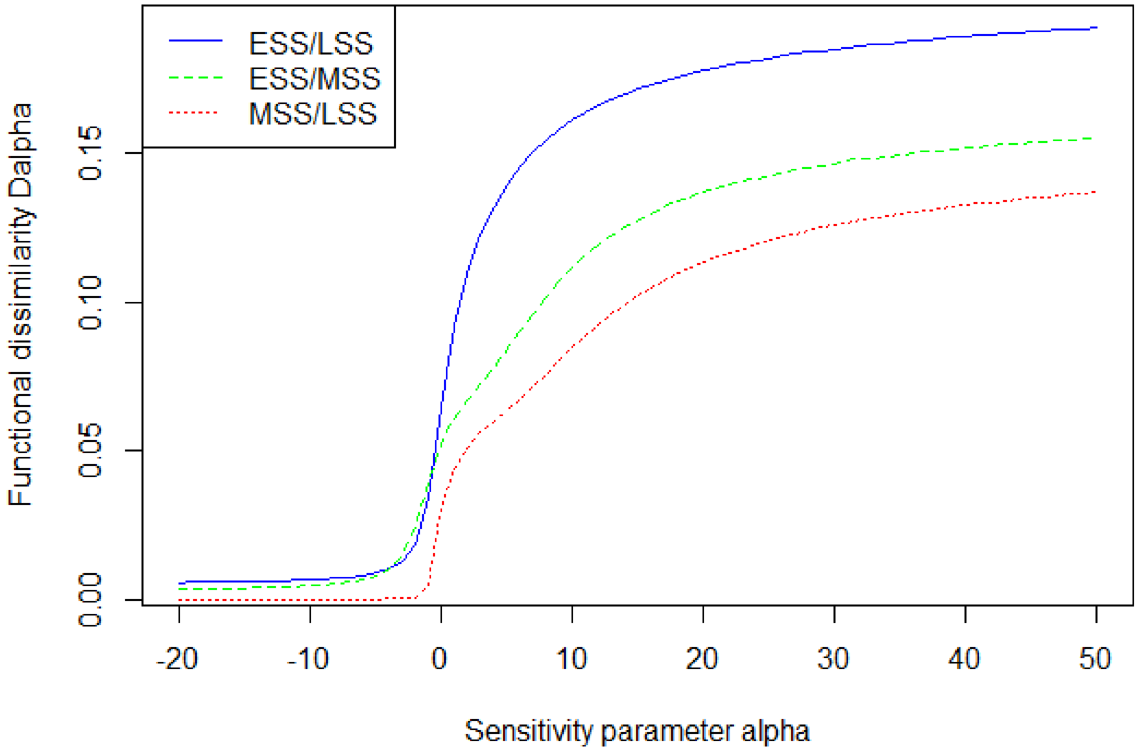


```
# The results above used the generalization of the Bray-Curtis index
proposed in the main text. The function generalized_Tradidiss_alpha
also allows to use the generalization Marczewski-Steinhaus
dissimilarity coefficient, and that of the evenness-based
dissimilarity coefficient both introduced in the main text (see
Manual of package adiv, function generalized_Tradidiss, for
details).
```

**References**


Pavoine, S. (2020) adiv: an R package to analyse biodiversity in ecology. Methods in Ecology and Evolution 11: 487-493. URL: https://doi.org/10.1111/2041-210X.13430.
