## Appendix 2 for "Towards a unifying framework for diversity and dissimilarity coefficients"

**Appendix 2.** Diversity partitioning of $Q_U^\alpha$.

Let $U$ and $V$ be two plots where $p_{Uj}$ is the relative abundance of species $j$ in plot $U$ (with $0 \le p_{Uj} \le 1$ and $\sum_{j=1}^{N} p_{Uj} = 1$), $N$ is the total number of species in both plots and $\mathbf{D} = [d_{ij}]$ is a quadratic dissimilarity matrix between species $i$ and $j$ in the range $[0,1]$ such that $d_{ij} = d_{ji}$ and $d_{jj} = 0$.

A well-known option mentioned by Champely and Chessel (2002) to generalize Rao's quadratic diversity consists in expressing parametric diversity as:

$$C_U^\alpha = \begin{cases} \sum_{i,j}^{N} p_{Ui} p_{Uj} d_{ij}^\alpha \text{ for } \alpha > 0 \\ C_U^0 = 1 - \sum_{j=1}^{N} p_{Uj}^2 \end{cases} \quad \text{(A1)}$$

If the interspecies dissimilarities $d_{ij}$ are squared Euclidean, $C_U^\alpha$ is concave for $0 < \alpha \le 1$, as for any squared Euclidean dissimilarity matrix $\mathbf{D} = [d_{ij}]$, the Hadamard power matrix $\mathbf{D}^{(\alpha)} = [d_{ij}^\alpha]$ is also squared Euclidean for $0 < \alpha \le 1$ (Schoenberg 1938; Rao 1984).

A matrix of dissimilarities $d_{ij}$ among $N$ species is said to be squared Euclidean, if the $N$ species can be embedded in a Euclidean space such that the Euclidean distance between species $i$ and $j$ is $\sqrt{d_{ij}}$ (Gower and Legendre 1986). Note that while Euclidean dissimilarities are automatically also squared Euclidean, the reverse is not true.

In addition, it is also true that, if the matrix $\mathbf{D}^{(\alpha_0)}$ is squared Euclidean for some $\alpha_0 > 0$, then for every $0 < \alpha \le \alpha_0$, the matrix $\mathbf{D}^{(\alpha)}$ is squared Euclidean and the parametric diversity index $C_U^\alpha = \sum_{i,j}^{N} p_{Ui} p_{Uj} d_{ij}^\alpha$ is concave on the set of $N$-dimensional distribution of species abundances (on the $N$-dimensional unit simplex).

In fact, if $\mathbf{D}^{(\alpha_0)}$ is squared Euclidean for $\alpha_0 > 0$, then $\mathbf{D}^{(\alpha)}$, $0 < \alpha \le \alpha_0$ is also squared Euclidean because $\frac{\alpha}{\alpha_0} \le 1$ and $\left(\left(d_{ij}^{\alpha_0}\right)^{\alpha/\alpha_0}\right) = \left(d_{ij}^\alpha\right) = \mathbf{D}^{(\alpha)}$.

From the concavity of $C_U^\alpha$ it follows that (Rao 1982):

$$\frac{1}{2}(C_U^\alpha + C_V^\alpha) \le C_{UV}^\alpha \quad \text{(A2)}$$

where $C_U^\alpha = \sum_{i,j}^{N} p_{Ui} p_{Uj} d_{ij}^\alpha$, $C_V^\alpha = \sum_{i,j}^{N} p_{Vi} p_{Vj} d_{ij}^\alpha$, and $C_{UV}^\alpha = \sum_{i,j}^{N} p_{Ui} p_{Vj} d_{ij}^\alpha$ is the expected dissimilarity between two individuals drawn at random, one from plot $U$ and one from plot $V$.

Let $\pi_{Uij} = p_{Ui} \times p_{Uj}$ be the joint probability of the pair of species $(i, j)$ in plot $U$ and $\Pi_{ij} = p_{Ui} \times p_{Vj}$ be the joint probability of the pair of species $(i, j)$ one from plot $U$ and one from plot $V$ in this order.

Since $\frac{1}{2}(C_U^\alpha + C_V^\alpha) = \frac{1}{2}(\sum_{i,j}^N p_{Ui} p_{Uj} d_{ij}^\alpha + \sum_{i,j}^N p_{Vi} p_{Vj} d_{ij}^\alpha) = \sum_{i,j}^N \frac{1}{2}(p_{Ui} p_{Uj} + p_{Vi} p_{Vj}) d_{ij}^\alpha$ , we have:

$$\sum_{i,j}^N \bar{\pi}_{ij} d_{ij}^\alpha \leq \sum_{i,j}^N \Pi_{ij} d_{ij}^\alpha \qquad \text{(A3)}$$

where $\bar{\pi}_{ij} = \frac{1}{2}(\pi_{Uij} + \pi_{Vij})$. Therefore, for $0 < \alpha \leq 1$ it follows that

$$\left(\sum_{i,j}^N \bar{\pi}_{ij} d_{ij}^\alpha\right)^{\frac{1}{\alpha}} \leq \left(\sum_{i,j}^N \Pi_{ij} d_{ij}^\alpha\right)^{\frac{1}{\alpha}} \qquad \text{(A4)}$$

Although the inequality in A4 does not represent the usual diversity partitioning into the classical alpha, beta and gamma components, nonetheless it provides an adequate approach to decompose the parametric diversity $Q_U^\alpha = \left(\sum_{i,j}^N p_{Ui} p_{Uj} d_{ij}^\alpha\right)^{\frac{1}{\alpha}}$ into a mean within-plot diversity $\bar{Q}_{UV}^\alpha = \left(\sum_{i,j}^N \bar{\pi}_{ij} d_{ij}^\alpha\right)^{\frac{1}{\alpha}}$ and a between-plot diversity $Q_{UV}^\alpha = \left(\sum_{i,j}^N \Pi_{ij} d_{ij}^\alpha\right)^{\frac{1}{\alpha}}$.

Both diversity components can be further used to derive a plot-to-plot dissimilarity coefficient that is based on the excess of between-plot diversity compared to mean within-plot diversity as:

$$\Theta_{UV}^\alpha = \frac{Q_{UV}^\alpha - \bar{Q}_{UV}^\alpha}{1 - \bar{Q}_{UV}^\alpha} \qquad \text{(A5)}$$

For $0 < \alpha \leq 1$, $\Theta_{UV}^\alpha$ thus represents a normalized diversity-derived dissimilarity coefficient in the range $[0,1]$ that is obtained by linearly rescaling between-plot diversity $Q_{UV}^\alpha$ in the usual way: $\left(Q_{UV}^\alpha - \min Q_{UV}^\alpha\right) / \left(\max Q_{UV}^\alpha - \min Q_{UV}^\alpha\right)$. For details, see Pavoine and Ricotta (2014).

From the above, it also follows that if the matrix $\mathbf{D}^{(\alpha_0)}$ is squared Euclidean for some $\alpha_0 > 0$, then for every $0 < \alpha < \alpha_0$, the coefficient $\Theta_{UV}^\alpha$ represents an adequate dissimilarity index in the range $[0,1]$.

**References**


Champely, S., Chessel, D. (2002) Measuring biological diversity using Euclidean metrics. Environmental and Ecological Statistics 9: 167–177.

Gower, J.C., Legendre, P. (1986) Metric and Euclidean properties of dissimilarity coefficients. Journal of Classification 3: 5–48.

Pavoine, S., Ricotta, C. (2014) Functional and phylogenetic similarity among communities. Methods in Ecology and Evolution 5: 666–675.

Rao, C.R. (1982) Diversity and dissimilarity measurements: A unified approach. Theoretical Population Biology 21: 24–43.

Rao, C.R. (1984) Convexity properties of entropy functions and analysis of diversity. In: Inequalities in Statistics and Probability (Y.L. Tong, Ed.), IMS Lecture Notes Monograph Series, Vol. 5, p. 68–77, Hayward, CA.

Schoenberg, I.J. (1938). Metric spaces and positive definite functions. Transactions of the American Mathematical Society 44: 522–536.
